# Antimicrobial Effects of Ibuprofen Combined with Standard of Care Antimicrobials against Cystic Fibrosis Pathogens

**DOI:** 10.1101/2021.05.11.443633

**Authors:** Qingquan Chen, Marleini Ilanga, Sabona B. Simbassa, Bhagath Chirra, Kush N. Shah, Carolyn L. Cannon

**Affiliations:** Department of Microbial Pathogenesis and Immunology, Texas A&M University Health Science Center, College Station, TX 77843, USA

## Abstract

Cystic Fibrosis (CF) is a common fatal genetic disease caused by mutations happened to cystic fibrosis transmembrane conductance regulator (CFTR) gene. Lungs of CF patients are often colonized or infected with microorganisms. Drug resistant bacterial infection has been problematic in cystic fibrosis patient. The chronic bacterial infections and concomitant airway inflammation could damage the lung and lead to respiratory failure. Several clinical trials have demonstrated that high-dose ibuprofen reduces the rate of pulmonary function decline in CF patients. This beneficial effect has been attributed to the anti-inflammatory properties of ibuprofen. Previously, we have confirmed that high-dose ibuprofen demonstrated antimicrobial activity against *P. aeruginosa* in *in vitro* and *in vivo*. However, no study has examined the antimicrobial effect of combining ibuprofen with standard-of-care (SoC) antimicrobials. Here, we evaluated possible synergistic activity of combinations of common nonsteroidal anti-inflammatory drugs (NSAIDs), namely, aspirin, naproxen, and ibuprofen, with commonly used antibiotics for CF patients. The drug combinations were screened against different CF clinical isolates. Drugs that demonstrated efficacy in the presence of ibuprofen were further verified synergistic effects between these antimicrobials and NSAIDs. Finally, the survival analysis of an *P. aeruginosa* murine infection model was used to demonstrate the efficacy of synergistic combination. Our results suggest that combinations of ibuprofen with commonly used antibiotics demonstrate synergistic antimicrobial activity against drug resistant, clinical bacterial strains in *in vitro*. The efficacy of combination ceftazidime and ibuprofen was demonstrated in *in vivo*.

## Introduction

Chronic infection and inflammation are the hallmarks of cystic fibrosis (CF) lung disease, which are responsible for majority of cases of morbidity and mortality in CF patients (1-3). Chronic infection elicits an acute inflammation response, which is characterized with persistent neutrophil influx(4). However, because of distinctly altered lung environment in CF patient, the inflammation fails to clear the infection, resulting worse airway obstruction, chronic endobronchial infection, and excessive airway inflammation(4, 5). Although clearance pulmonary obstruction and bacterial infection is the main focus of CF therapy, because of the effects of inflammation on lung destruction, more and more studies investigated on therapies against excessive inflammatory response(4, 6, 7). Several in vivo models and clinical trials demonstrated appreciable clinical effect of oral and inhaled corticosteroids, macrolides, and nonsteroidal anti-inflammatory drugs (NSAIDs), such as ibuprofen(8-13).

Because of its effectiveness and safety profile, ibuprofen is advantageous as an anti-inflammatory drug. Compared to corticosteroid that causes growth retardation and cataracts, ibuprofen has less safety concerns, mainly with GI upsets. Ibuprofen has demonstrated to reduce the recruitment of neutrophil into the airway in both a mouse model of acute *Pseudomonas* pulmonary infection and a rat model of endotoxin-induced alveolitis(14, 15). In a chronic Pseudomonas endobronchial infection study, rat treated orally with ibuprofen, resulting in a drug plasma concentration of 55+24 µg/L, achieved a significant reduction in the inflammatory response and improved weight gain compared to control treatment(16). At the same concentration, ibuprofen reduced the level of leukotriene B4 production without reducing pulmonary bacterial burden(16). Furthermore, Konstan et al. conducted a randomized, double-blind, placebo-controlled clinical trial to evaluate the safety and efficacy of high-dose ibuprofen (50 to 100 µg/mL) in CF patients. High-dose ibuprofen reduced the rate of decline of pulmonary function in CF patient with good to excellent pulmonary function(10). Then, patients, between 6 to 17 years old with a baseline forced expiratory volume in 1 s (FEV1) of >60%, demonstrated a significantly lower rate of decline of the FEV1 percentage predicted when treated with high-dose ibuprofen than patients treated with placebo(17). Lands et al. also investigated the safety and efficacy of high-dose ibuprofen in CF children between 6 to 18 years old. This randomized, multicenter, double-blinded, placebo-controlled trial did not show statistically significant difference in the mean annual rate of decline in FEV1. However, the annual rate of decline of the forced vital capacity (FVC) percentage predicted significantly decreased in patients treated with high-dose ibuprofen(8). To sum up, these clinical trials suggested the benefits and relative safety of long-term use of high-dose ibuprofen in CF patients as well as attribute these benefits to the anti-inflammatory property of ibuprofen.

Studies have documented the antimicrobial and antifungal activity of ibuprofen. Our lab demonstrated the direct effect of ibuprofen on bacterial pathogens prevalent in the CF lung. Shah et al. described the effects of high-dose ibuprofen on two important Gram-negative bacterial pathogens responsible for chronic pulmonary infection in CF patients, *Pseudomonas aeruginosa* in *in vitro* and *in vivo* studies(18). The results demonstrated ibuprofen has antimicrobial effects against laboratory and clinical isolates of both *P. aeruginosa* and *Burkholderia* spp. in a dose-dependent manner. In an acute murine pneumonia model, mice orally administrated with high-dose ibuprofen showed a reduced bacterial burden in lung, and superior survival compared to mice with sham treatment(18). Studies have demonstrated that several NSAIDs, including ibuprofen, and aspirin, are synergistic with cefuroxime and chloramphenicol against MRSA(19). We hypothesized that the remarkable results of the CF ibuprofen trials illustrate that, in addition to its anti-inflammatory properties, ibuprofen prevents lung function decline through inhibition of bacterial growth. Hence, we propose that ibuprofen and other NSAIDs, such as naproxen and aspirin, sensitize drug-resistant bacteria to established antimicrobials, thus exerting a synergistic bactericidal effect. Ibuprofen and possibly other NSAIDs may prove to be ideal agents with dual anti-inflammatory and antimicrobial activity that may be of great treatment benefit to the CF patient without incurring the attendant risks. Developing such effective combinations comprised of therapeutics already approved for human use will also allow us to rapidly devise novel, effective intervention strategies for the treatment of chronic lung infections with multi-drug resistant pathogens found in the lungs of CF patients.

## Result

### The zone of inhibition of antibiotic-infused disc was determined against CF clinical isolates

We characterized the zone of inhibition of antibiotic-infused disc against 1 *P. aeruginosa* CF clinical isolates (PA HP3), 1 *E. meningoseptica* (EM) CF isolates (EM 2-18), 1 *A. xylosoxidans* (AX) CF isolates (AX 2-79), 1 *H. influenzae* (HI) CF isolates (HI 2501), and 1 MRSA CF clinical isolates (SA LL06) (**Table 1**). The Antimicrobial susceptibility of each drug against CF clinical isolates was determined according to the clinical laboratory standard breakpoints. The green indicates bacteria is sensitive to the tested antimicrobial. The yellow indicates bacteria is intermediate resistant to the tested antimicrobials. The red indicates bacteria is resistant to the tested antimicrobials. All isolates demonstrated resistance to at least 2 tested antibiotics. Particularly, 2 strains indicated in orange color were multi-drug resistant. After supplement with high-dose ibuprofen, aztreonam and ceftazidime showed significant zone of inhibition against PA HP3, and amikacin showed significant increase in zone of inhibition against EA 2-18 (**Figure 1**). Amikacin demonstrated a significant zone of inhibition against AX 2-79 with the addition of high-dose ibuprofen (**Figure 1**). Ceftazidime demonstrated significant increase in zone of inhibition against HI 2501 after adding high-dose ibuprofen. Gentamicin and vancomycin with addition of high-dose ibuprofen showed significant increase in zone of inhibition against SA LL06 (**Figure 1**).

**Table 1.** The zone of inhibition of antibiotic-infused disc against Cystic Fibrosis pathogens.

| Strains | Average zone of inhibition (unit:mm) |  |  |  |  |  |  |  |
| --- | --- | --- | --- | --- | --- | --- | --- | --- |
|  | AMK | ATM | CAZ | CST | TOB | GEN | LVX | VAN |
| PA HP3 | 19 | 18 | 13 | 16 | 6 | - | - | - |
| EM 2-18 | 11 | 6 | 6 | 6 | 6 | - | - | - |
| AX 2-79 | 8 | 7 | 23 | 9 | 6 | - | - | - |
| HI 2501 | 14 | 24 | 24 | 14 | 15 | - | - | - |
| SA LL06 | - | - | - | - | - | 22 | 9 | 14 |

**Figure 1.**
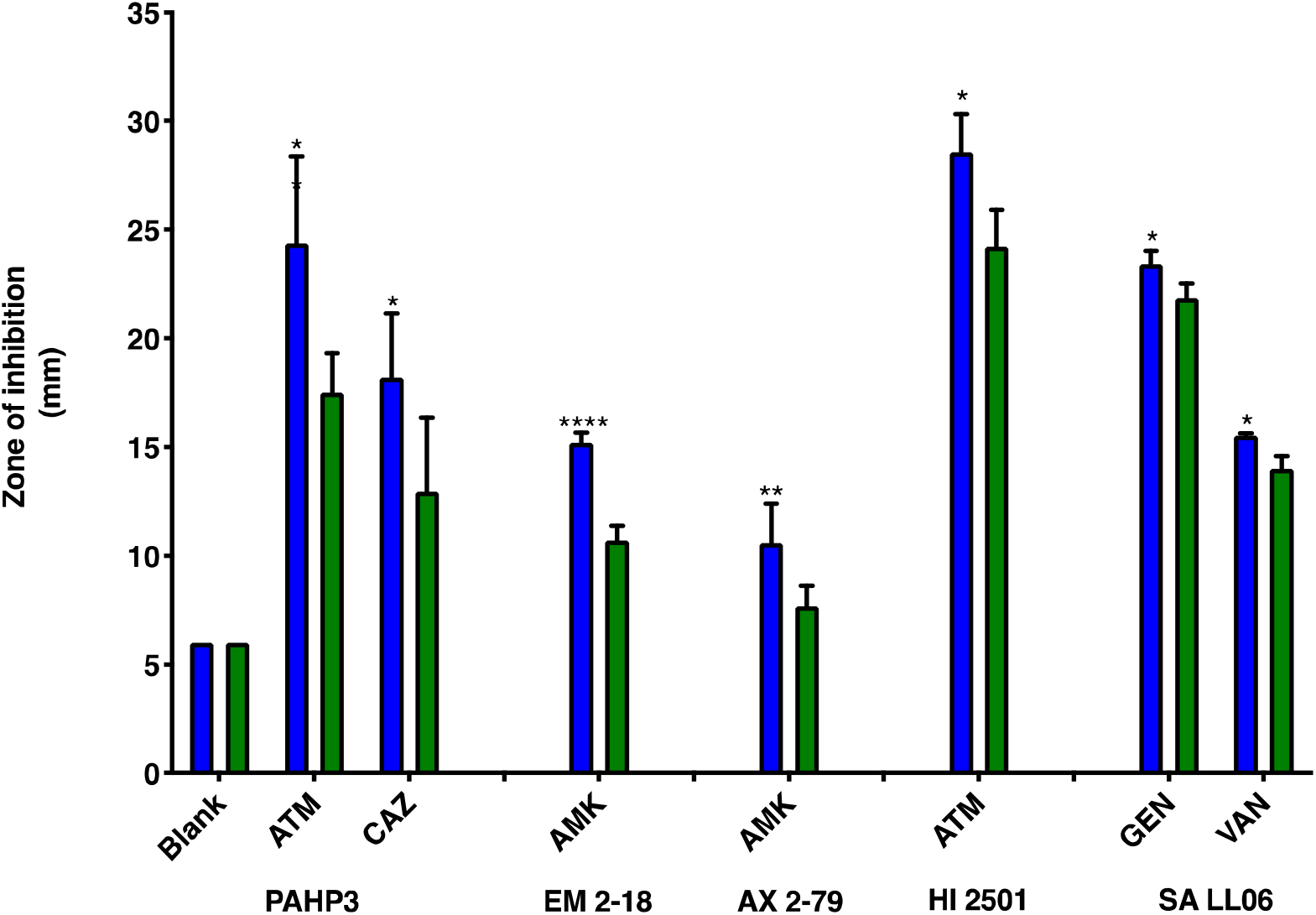
Supplementing ibuprofen improved the zone of inhibition of antibiotics-infused disc against tested CF clinical isolates. Statistical significance determined by two-way ANOVA followed by Tukey’s multiple comparison test (* indicates p≤0.05, ** indicates p≤0.01, and **** indicates p≤0.0001).

### *In vitro* antimicrobial activity of ibuprofen, naproxen, aspirin, amikacin, aztreonam, and ceftazidime against *P. aeruginosa* and *E. meningoseptica*

We characterized the minimum inhibitory concentration (MIC) of ibuprofen, naproxen, aspirin against PA HP3 and EM 2-18 (**Table 2)**, amikacin against EM 2-18, and aztreonam and ceftazidime against PA HP3 (**Table 3**). Ibuprofen demonstrated mild antimicrobial activity, which the MIC is 512 µg/mL against PA HP3 and 256 µg/mL against EM 2-18, which consistent with our previous observation showed that ibuprofen has mild antimicrobial activity against *P. aeruginosa* laboratory strain and clinical isolates. The MIC of naproxen is 2048 µg/mL against EM 2-18. However, we were not be able to detect the MIC of naproxen against PA HP3 and MIC of aspirin against PA HP3 and EM 2-18 within our concentration upper limit, which was 2048 µg/mL. Other studies have observed that the MIC of naproxen and aspirin are above 3 mg/mL against gram-negative bacteria, which is above our designed assay detection upper limit. The MIC of aztreonam and ceftazidime against PA HP3 is 4 and 16 µg/mL, respectively. The MIC of amikacin against EM 2-18 is 16 µg/mL.

**Table 2.** The minimum inhibitory concentration of ibuprofen, naproxen, and aspirin against PA HP3 and PA 2-18.

|  | **PA HP3** | **EM 2-18** |
| --- | --- | --- |
| **Ibuprofen** | 512 | 256 |
| **Naproxen** | >1024 | 2048 |
| **Aspirin** | >1024 | >1024 |

**Table 3.** The MIC of aztreonam and ceftazidime against PA HP3, and amikacin against EM 2-18 as well as combining antibiotics with NSAIDs against selected isolates of *Pseudomonas aeruginosa* and *Elizabethkingia meningoseptica*.

|  | MIC µg/mL | MIC ([Ibuprofen]) µg/mL | MIC ([Naproxen]) µg/mL | MIC ([Aspirin]) µg/mL |
| --- | --- | --- | --- | --- |
| **Aztreonam (PA HP3)** | **4** | **2 (50)** | **2 (75)** | **4 (100)** |
| **Ceftazidime (PA HP3)** | **16** | **4 (50)** | **12 (50)** | **16 (100)** |
| **Amikacin (EM 2-18)** | **16** | **12 (50)** | **12 (100)** | **16 (100)** |

### Synergistic effect of ibuprofen and ceftazidime against PA HP3 demonstrated by checkerboard assay

To explore the potential synergistic antimicrobial effects between antibiotics and NSAIDs, we tested combined drug against PA HP3 and EM 2-18 using a checkerboard assay. The MICs of antibiotics were reduced in **Table 3** with the presence of 50 µg/mL ibuprofen and various concentrations of naproxen. However, the MIC of antibiotics did not change with the presence of highest concentration of aspirin. Then, we used the fractional inhibitory concentration to interpret potential drug combination effects. Based on the FIC calculation performed (citation), we determined that ceftazidime is synergistic with ibuprofen against PA HP3. Ibuprofen is additive with aztreonam against PA HP3 and with amikacin against EM 2-18. Naproxen is additive with all three antibiotics (**Table 3**). Furthermore, the combinational MICs are determined based on turbidity of liquid assay in 96-well plates, which has poor sensitivity and is the limitation of checkerboard assay. Hence, we decide to use 24-hour endpoint CFU study to further examine the potential synergistic drug combination.

### Synergistic effects of NSAIDs and antibiotics demonstrated by endpoint CFU studies

Even though we observe a synergistic effect between ceftazidime and ibuprofen against PA HP3 using a checkerboard assay, we further performed an endpoint CFU study to investigate the effect of the combination therapeutic on CFU. The concentrations used in 24-hour endpoint CFU study were selected based on the checkerboard assay result. For 24-hour end-point CFU study, we selected NSAIDs and antibiotics concentrations at sub or at individual MIC concentrations but including the combinational MIC within the testing range. The bacterial concentration of PA HP3 is ∼10^9^ CFU/mL when treated with 2 μg/mL aztreonam alone. However, following supplementing of 100 μg/mL naproxen, the bacterial burden of PA HP3 is reduced to ∼10^6^ CFU/mL (**Figure 2A**). Since the synergistic effect in endpoint CFU study is defined as a ≥ 2-log_10_ reduction in bacterial burden compared with the most efficacious individual treatment, the aforementioned combination of aztreonam and naproxen demonstrated synergy in our endpoint CFU study. When we treated PA HP3 with 8 μg/mL ceftazidime alone, the bacterial concentration of PA HP3 is ∼10^7^ CFU/mL. When we added 100 μg/mL naproxen, the bacterial burden was reduced to ∼10^4^ CFU/mL, which indicated synergy (**Figure 2B**). Furthermore, we verified that ibuprofen was synergistic with all three antibiotics. With addition of 100 μg/mL ibuprofen, 1 μg/mL aztreonam achieved ∼ 6-log_10_ CFU/mL reduction compared to individual treatment (**Figure 3A**); 2 μg/mL ceftazidime achieved ∼4-log_10_ bacterial burden reduction compared to individual treatment (**Figure 3B**); 4 μg/mL amikacin achieved ∼3-log_10_ reduction compared to individual treatment (**Figure 3C**). Both naproxen and ibuprofen demonstrated synergistic in combination with aztreonam, ceftazidime, and amikacin. Furthermore, ibuprofen demonstrated a greater reduction of bacterial burden when combined with all three antibiotics. The high-dose ibuprofen has been used in CF patients as anti-inflammatory drug. Hence, we decide to process to test the efficacy of antibiotics combined with ibuprofen in murine pneumonia model.

**Figure 2.**
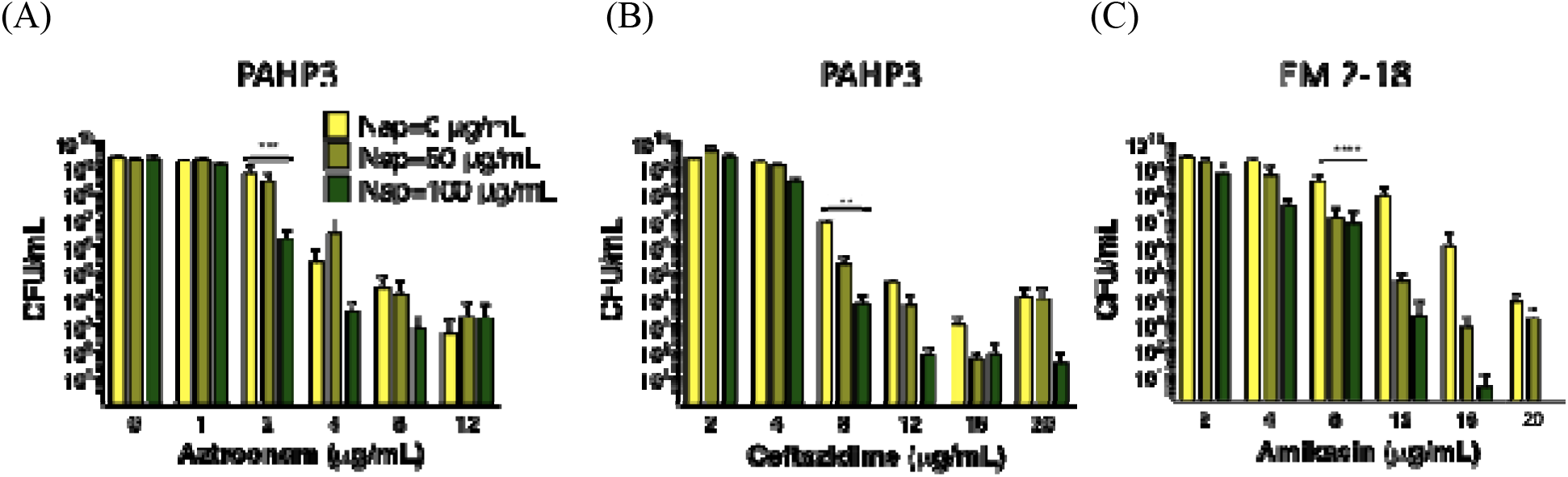
Synergy demonstrated between naproxen and (A) aztreonam, (B) ceftazidime, and (C) amikacin against PA HP3 and EM 2-18 by endpoint CFU study after 24-hour incubation with the drug concentration ratios (in mg/mL) indicated under each panel. Data are shown as mean and standard deviation (n = 6). Statistical significance determined by one-way ANOVA followed by Tukey’s multiple comparison test (** indicates p≤0.01, *** indicates p≤0.001, and **** indicates p≤0.0001).

**Figure 3.**
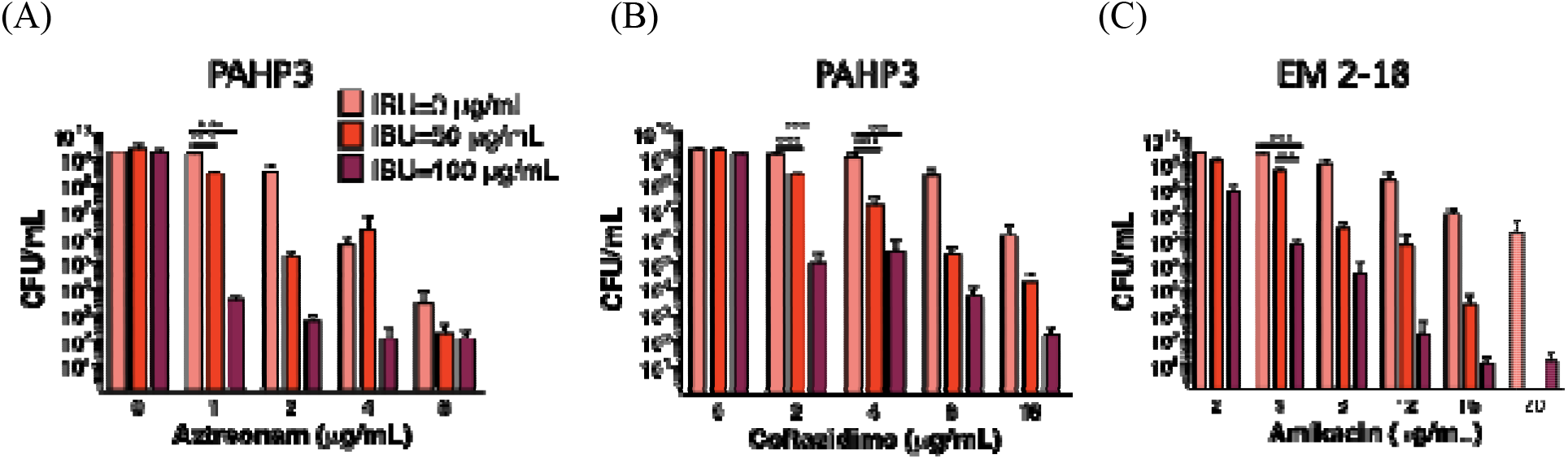
Synergy demonstrated between ibuprofen and (A) aztreonam, (B) ceftazidime, and (C) amikacin against PA HP3 and EM 2-18 by endpoint CFU study after 24-hour incubation with the drug concentration ratios (in mg/mL) indicated under each panel. Data are shown as mean and standard deviation (n = 6). Statistical significance determined by one-way ANOVA followed by Tukey’s multiple comparison test (** indicates p≤0.01, *** indicates p≤0.001, and **** indicates p≤0.0001).

### Ibuprofen in combination with ceftazidime improved mice survival significantly an acute pneumonia infection

To investigate the significance of combinational therapy in a murine acute pneumonia infectious model, mice were intranasally infected with PA HP3 and treated with control, ceftazidime via intraperitoneal injection only, ibuprofen via oral feeding only, and ceftazidime via intraperitoneal injection combined with ibuprofen via oral feeding and monitored host health and survive. Infected mice were treated every 8 hours up to 7 doses. The experiment last 72 hours. At 72 hours, infected mice treated with combinational therapy demonstrated a significant survival advantage over individually treated group of mice (**Figure 4**).

**Figure 4.**
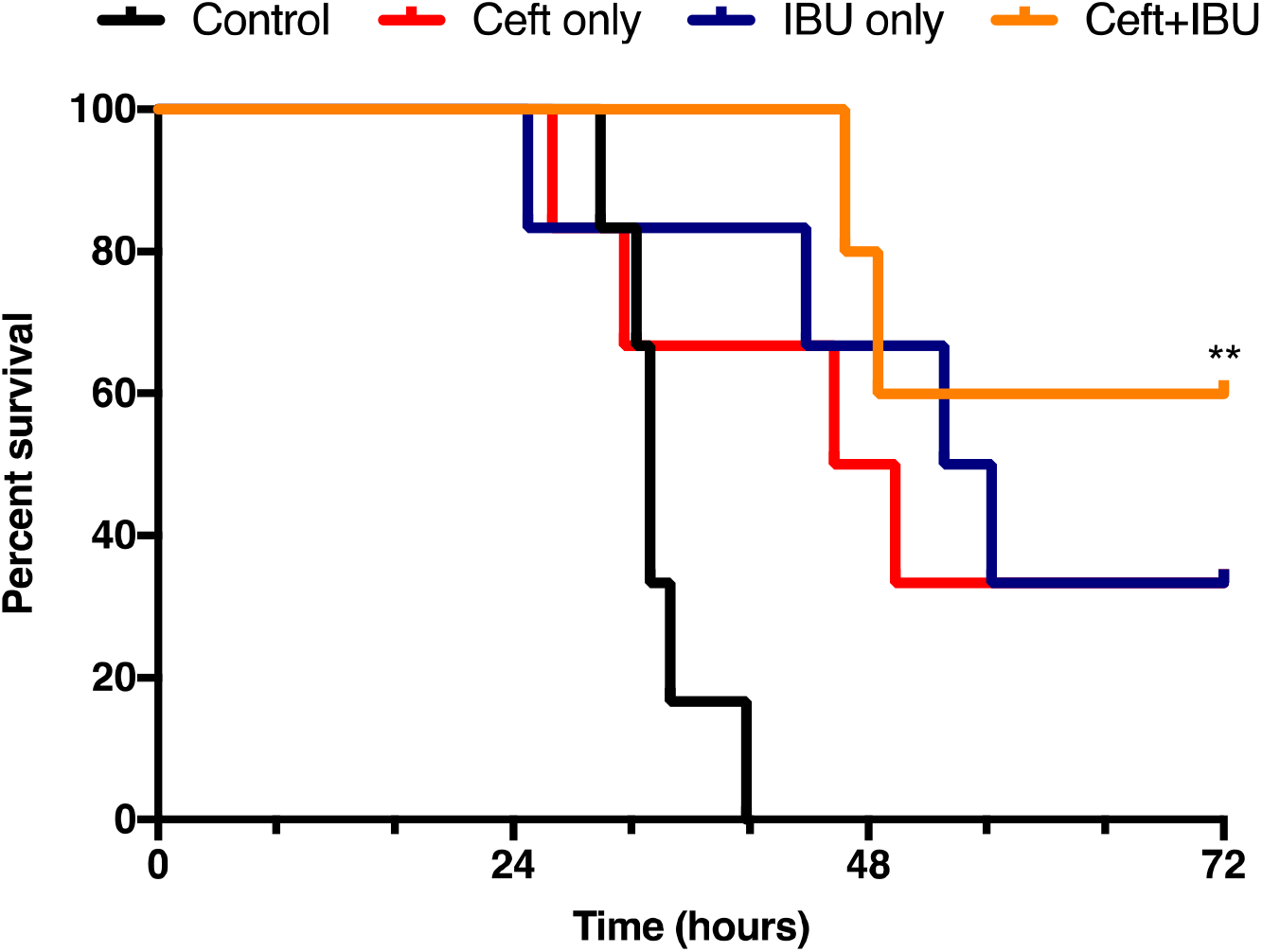
The combinational therapy of ibuprofen and ceftazidime demonstrated a significantly survival advantage in a murine pneumonia infectious model (n=6). Statistical significance determined by Mantel-Cox test (** indicates p≤0.01).

## Discussion

The disc diffusion assay is implemented for its rapid results turnover rate. Commercially available drug contained discs were used in the test. Studies have suggested that disc diffusion might be insensitive as well inaccurate. However, this assay provides a rapid screening means to identify the drug combinations that ibuprofen could enhance the antimicrobial activity of commonly used antibiotics.

Other studies, including our own observation, have demonstrated that NSAIDs are synergistic with antibiotics against gram-negative and gram-positive pathogens(19).

To sum up, in combination with standard of care antimicrobials, we were able to demonstrate that ibuprofen could significantly increase the zone of inhibition against CF clinical isolates. The drug combination obtained in disc diffusion assay provides us candidates to proceed for detail microbiological characterization. *In vitro* studies suggested that ibuprofen has antimicrobial activity against CF clinical isolates, which confirmed by our previous observation. Despite aspirin did not demonstrate either synergy or additive effects with selected antibiotics, naproxen and ibuprofen showed additive effects and synergistic effects with selected antibiotics. Further, the 24 hour endpoint CFU study confirmed that naproxen and ibuprofen are synergistic with either one of three antibiotics. Ibuprofen in combined with antibiotics exerted greater reduction of bacterial burden. Finally, when mice treated with sub-MIC concentration of ceftazidime intraperitoneally and ibuprofen orally, infected mice demonstrated improved survival rates compare to single drug treated mice. Our i*n vitro* and *in vivo* experiments suggests that ibuprofen has mild antimicrobial activity in addition to it anti-inflammatory property, particularly after combining with standard of care antibiotics. Currently, high-does ibuprofen treatment is not widely available in CF centers. It would be worth revisiting ibuprofen to explore its clinical benefits in CF patients.

## Method

### Bacterial strains

All CF strains of *Pseudomonas aeruginosa, Achromobacter xylosoxidans, Burkholderia* spp., *Elizabethkingia meningoseptica, Haemophilus influenzae*, MRSA, and *Stenotrophomonas maltophilia* were utilized for our studies and comprised laboratory as well as clinical isolates. *Pseudomonas aeruginosa* laboratory strain PAO1 was generously donated by Gerald Pier (Harvard University, Boston, MA). The CF mucoid clinical isolates *P. aeruginosa* M57-15 was graciously donated by Thomas Ferkol (Washington University, St. Louis, MO) (84 Parth paper). The remaining *P. aeruginosa* CF clinical isolates (PA LF05, PA HP3, PA 2-9, PA 2-15, PA 2-22, PA 2-23, PA 2-26, PA 2-45, PA 2-51, and PA 3-39) were isolated from the sputa of cystic fibrosis patients at St. Louis Children’s Hospital. *Achromobacter xylosoxidans* CF clinical isolates (AX 2-79 and AX 3-26), *Elizabethkingia meningoseptica* CF clinical isolates (EM 2-14 and EM 2-18), MRSA (SA EH05 and SA LL06), and *Stenotrophomonas maltophilia* (SM AH06 and SM AH08) were isolated from the sputa of CF patients at St. Louis Children’s Hospital. The *Burkholderia* spp., including *Burkholderia cenocepacia* J2315, *burkholderia dolosa* (CF clinical isolates BD-F, and BD-G), *burkholderia gladioli* (CF clinical isolate BG 5291), and *burkholderia multivorans* (CF clinical isolate BM 2-6) were isolated from the sputa of CF patients at St. Louis Children’s Hospital. *Haemophilus influenzae* (HI 4315, HI 2501, and HI 3864) were kindly provided by Joseph St. Geme (University of Pennsylvania, Philadelphia, PA).

### Bacteria culture

Bacteria were streaked from frozen glycerol stock onto tryptic soy agar (TSA, BD BBL) or chocolate agar (Hardy Diagnostics) plates and incubated overnight at 37 °C with 5% carbon dioxide (CO_2_) until individual colonies formed. A single colony was inoculated into 5 mL Miller Hinton (MH, BD Difco) or Brain Heart Infusion (BHI, BD Difco) media and grown at 37 °C in a shaking incubator at 200 rpm to an OD_650_ of 0.4, which corresponds to ∼5 ⍰10^8^ CFU/mL. Bacteria cultures were adjusted to 5 ⍰10^8^ CFU/mL to prepare a working stock for all experiments.

### Disc diffusion assay

MH agar plates were prepared by autoclaving MH with 17g agar per liter of media. After autoclaving, the agar was cooled to 70 °C and 100 μg/mL solution of ibuprofen dissolved in DMSO was added to a final concentration of 100 μg/mL. An equivalent amount of DMSO was added to another batch of MH agar to serve as control. Plates were cast from the ibuprofen added MH agar and DMSO added MH agar. 100 μL of bacteria culture were dispensed onto agar plates and spread evenly, after which antibiotic (amikacin, aztreonam, ceftazidime, colistin, and tobramycin)-infused discs were placed on top of the agar. Plates were incubated between 18-24 hours. Susceptibility was determined by measuring the diameter of the zone of growth inhibition.

### *In vitro* antimicrobial activity

Minimum inhibitory concentrations (MIC) were determined according to the standard Clinical and Laboratory Standards Institute (CLSI) broth-microdilution method and adapted from previous (minocycline paper). Briefly, 100 μL working stock of bacterial suspension was added to each well (n=3) containing 100 μL ibuprofen, naproxen, aspirin, ceftazidime, amikacin, or aztreonam solution in a 96 well plate. All solutions were comprised of 95% MH broth and 5% (v/v) DMSO. Bacteria were incubated with 0.06, 0.13, 0.25, 0.5, 1, 2, 4, 8, 16, 32, 64, 128 μg/mL amikacin, aztreonam, or ceftazidime, or 1, 2, 4, 8, 16, 32, 64, 128, 256, 512, and 1024 μg/ml ibuprofen, naproxen, or aspirin at 37 °C for 18 -24 hours under static conditions. The final concentration of DMSO in the assay was 2.5% (v/v). The MIC was determined as the lowest concentration that did not show any signs of bacterial growth upon visual inspection. All experiments were performed in triplicate.

### Determination of synergistic drug combinations

One *P. aeruginosa* (PA HP3) and one *E. meningoseptica* (EM 2-18) isolates were selected based on their susceptibility in disc diffusion assay. The final drug concentrations of ibuprofen, naproxen, and aspirin were 0, 50, 75, and 100 μg/ml. Based on the MIC values, a dynamic concentration scale for amikacin, aztreonam, and ceftazidime was used to determine the optimal ratio of synergistic concentrations between the two therapeutic agents. The final drug concentrations of amikacin against EM 2-18 were 1, 2, 4, 8, 12, and 16 μg/ml. The final drug concentrations of aztreonam against PA HP3 were 0.25, 0.5, 1, 2, and 4 μg/ml. The final drug concentration of ceftazidime against PA HP3 were 1, 2, 4, 8, 12, and 16 μg/ml. The final solutions were comprised of 95% MH broth and 5% DMSO. A 100 μL working stock of bacterial suspension was incubated with a 100 μL solution of therapeutic agents (n = 3) for 18 hours at 37 °C. Wells demonstrating bacterial growth inhibition were identified visually to determine a synergistic MIC. All experiments were performed in duplicate. To evaluate for potential synergy, the fractional inhibitory concentration (FIC) was calculated as shown in **Equation 1** and defined in **Table 1**.

### Determination of bacterial burden for synergistic drug combinations

Potential synergy between combinations of amikacin, aztreonam, or ceftazidime and ibuprofen, or naproxen against *P. aeruginosa* and *E. meningoseptica* isolates PA HP3 and EM 2-18 at a final concentration of 10^6^ CFU/mL were determined using a 24-hour end point CFU study performed in triplicate. The concentrations of ibuprofen and naproxen tested against PA HP3 and EM 2-18 were 0, 50, 100 μg/mL. The concentrations of aztreonam and ceftazidime in combination with ibuprofen against PA HP3 were 0, 1, 2, 4, and 8 or 0, 2, 4, 8, and 16 μg/mL, respectively. The concentrations of amikacin in combination with ibuprofen tested against EM 2-18 were 2, 4, 8, 12, 16, and 20 μg/mL. The concentrations of aztreonam and ceftazidime in combination with naproxen against PA HP3 were 0, 1, 2, 4, 8, and 12 or 2, 4, 8, 12, 16, and 20 μg/mL, respectively. The tested concentrations of amikacin in combined naproxen against EM 2-18 were 2, 4, 8, 12, 16, and 20 μg/mL. Synergy was defined as ≥2-log_10_ CFU/mL reduction between combined agents and the most effective individual agent at 24 hours^46^. A 100 μL working stock of bacterial suspension was incubated with 100 μL drug solution (n = 3) in each well of a 96 well plate at 37 °C for 24 hours with constant shaking at 100 RPM. The final solutions were comprised of 97.5% MH broth and 2.5% (v/v) DMSO. Finally, a 10-fold serial dilution was performed in MH broth with the bacterial suspension from each well and 50 μL of each dilution was plated onto a blood agar plate. Plates were incubated for 18 hours and colonies counted to determine the CFU for each condition. The potential synergistic effects were determined as described above. All experiments were performed in duplicate.

### Acute Murine-*P. aeruginosa* lung infection model

Male C57BL/6J mice (The Jackson Laboratory, Bar Harbor, ME) aged 5 weeks were used for all acute lung infection studies [34, 35], which were approved by the Texas A&M University Institutional Animal Care and Use Committee (IACUC). Mice were weighed and randomly assigned into four groups and were housed in a barrier facility under pathogen-free conditions until bacterial inoculation. When necessary, animals were euthanized with an overdose of ketamine:xylazine followed by cardiac puncture for exsanguination, a method approved by our IACUC (TAMU) and are consistent with the recommendations of the Panel on Euthanasia of the American Veterinary Medical Association.

*P. aeruginosa* PA HP3 was grown in LB (LB), pelleted, washed with phosphate buffered saline (PBS), and resuspended to an OD_650_ of 2.4 in LB (corresponding ∼1.3 × 10^10^ CFU/mL, as determined by serial dilution and plating). Following anesthesia via intraperitoneal injection of ketamine (60 mg/kg) and xylazine (8 mg/kg) cocktail, mice were intranasally inoculated with 75 mL of bacteria inoculum in LB broth at an LD_100_ of ∼1 × 10^9^ CFU per mouse. To test the efficacy of combinational therapy against PA HP3, at 2 h post infection, mice were treated every 8 hours subsequently for a maximum of 7 doses. Controlled mice were intraperitoneally injected with 50 µL saline in water and orally administered with 50 µL of 50:50 strawberry syrup : water mix. Ibuprofen treated mice were intraperitoneally injected with 50 µL saline in water and orally administrated with 50 µL of 50:50 strawberry syrup : water ibuprofen suspension mix (1.5 mg ibuprofen). Ceftazidime treated mice were intraperitoneally injected with 50 µL of 10 mg/mL ceftazidime and orally administered with 50 µL of 50:50 strawberry syrup : water mix. Combination treated mice were intraperitoneally injected with 50 µL 10 mg/mL ceftazidime and orally administered with 50 µL of 50:50 strawberry syrup : water ibuprofen suspension mix (1.5 mg ibuprofen). Infected mice were weighed and assigned a clinical infection score [36, 37] every 24 hours post infection. Clinical score is a semi-quantitative metric that evaluates for signs of infection, and ranges from asymptomatic (score 0) to moribund (score 6) based on the resting posture (0-2), fur (0-1), and activity level (0-3) of the infected mice. Mice survival were monitored up to 72 hours.

### Statistical analysis

All statistics were calculated using JMP pro 13 for Macintosh (SAS Institute, Cary, North Carolina, USA, www.jmp.com). Differences between the treatments were investigated by one-way ANOVA followed by Tukey’s multiple comparison test (95% confidence intervals). * indicates p≤0.05, ** indicates p≤0.01, *** indicates p≤0.001, and **** indicates p≤0.0001. The *in vivo* survival curves in the infection model were compared using a Log-rank Mantel-Cox test; whereas, differences in weight change and clinical scores of mice receiving different treatments were analyzed using Kruskal-Wallis one-way ANOVA and Dunn’s multiple comparison test. Data were deemed to be significantly different for p ≤ 0.05.

